# Limited Water Stress Triggers Adaptive Leaf Trait Response in Australian Commercial Rice Without Compromising Grain Yield and Quality

**DOI:** 10.1101/2025.05.21.655428

**Authors:** Yvonne Fernando, Mark Adams, Markus Kuhlmann, Vito Butardo

**Affiliations:** Department of Chemistry and Biotechnology, Swinburne University of Technology, Hawthorn, Australia; Institute of Plant Genetics and Crop Plant Research (IPK), Germany

**Keywords:** water use efficiency, vegetative stage, stomatal leaf traits, non-stomatal leaf traits, grain quality, rice, drought tolerance, photosynthetic efficiency, cuticular wax, milling recovery

## Abstract

Water scarcity poses significant challenge for the Australian rice industry, driving the need for varieties with improved water use efficiency (WUE). This study evaluated 18 Australian elite japonica, two indica rice lines, and the West African *japonica* cultivar Moroberekan to assess stomatal and non-stomatal leaf traits contributing to WUE. Plants were cultivated under controlled glasshouse conditions with two water treatments: ponded (PW) and limited water (LW) maintained at 60-65% field capacity throughout the vegetative stage. Under LW, plants exhibited decreased photosynthetic efficiency, stomatal conductance, and stomatal density but increased cuticular and epicuticular wax (CEW) deposition, which helped conserve water. The rice cultivar Sherpa exhibited reduced sensitivity to regulation of gas exchange, stomatal adjustments and photosynthetic efficiency, and the greatest CEW deposition among the japonica varieties, indicating potential for improved WUE. Thousand grain weight showed no significant differences between the two water treatments. Brown rice yield and milling recovery were also unaffected. Spectral analysis of dehulled seeds suggested that LW stress increased protein content but had no effect on lipids and starch. These findings demonstrate that LW during the vegetative stage can induce water-conserving traits without compromising yield or grain quality; providing valuable insights for developing water-efficient rice varieties.

**Graphical abstract:** 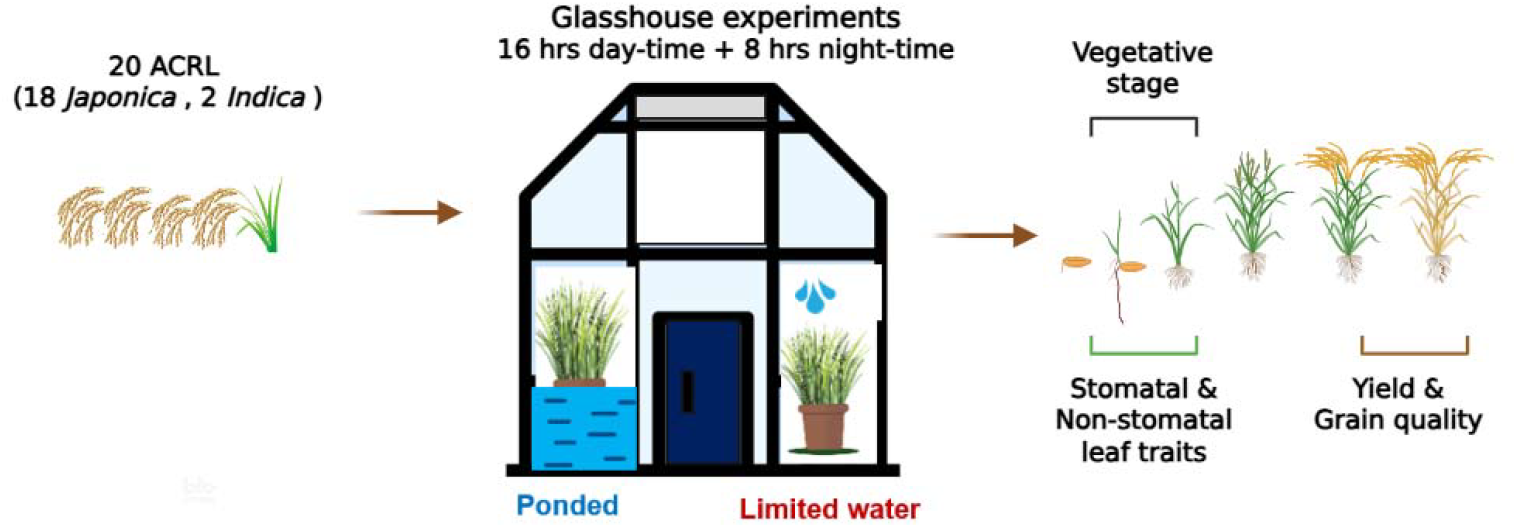

**Highlight:** Water scarcity elicits a range of stomatal and non-stomatal responses in Australian commercial rice varieties without compromising yield and grain quality while enhancing water conservation.

## Introduction

Water scarcity is an increasingly urgent global issue, particularly in regions where agriculture is a major economic driver (FAO, 2019; Rosa et al., 2020). Although the Australian rice industry is relatively small, it plays a significant role in the agricultural sector and contributes substantially to the economy (Bajwa & Chauhan, 2017; Department of Agriculture, 2022). While the Australian rice industry is amongst the most water-efficient globally (RGA, 2023), it nonetheless faces substantial challenges due to fluctuating water availability driven by variability in climate and in competing demands from other sectors. Water scarcity threatens food security because rice is a staple food for billions of people globally (Haonan et al., 2023; Hoque et al., 2021; Humphreys et al., 2006).

Improving water use efficiency (WUE) can help mitigate the impact of water scarcity on rice cultivation (Haonan et al., 2023; Mallareddy et al., 2023). Understanding mechanisms that influence WUE, particularly factors related to water loss through leaves, is essential to breeding and selecting water-efficient rice germplasm (Bramley et al., 2013; Lang et al., 2024). Leaf water loss via stomatal and non-stomatal pathways influences key traits including stomatal conductance and density, as well as cuticular and epicuticular wax deposition that collectively regulate plant water conservation (Pitaloka et al., 2022; Pitaloka et al., 2021).

Stomatal traits, including density, size, and arrangement, directly influence gas exchange and transpiration. Recent research has demonstrated that rice varieties with reduced stomatal density and smaller stomatal size exhibit improved WUE and biomass production under various growing conditions (Phunthong et al., 2024; Pitaloka et al., 2022). Non-stomatal traits such as cuticular wax composition and deposition also contribute substantially to plant water conservation, particularly under water-limited conditions (Haque et al., 1992; Srinivasan et al., 2008). Epicuticular waxes form a hydrophobic barrier on the leaf surface that reduces non-stomatal water loss while protecting against environmental stresses (Islam et al., 2009; Qin et al., 2011).

Most studies on WUE in rice have primarily focused on inducing severe drought conditions (Kamoshita et al., 2015; Praba et al., 2009), often resulting in reduced yield and grain quality (Blakeney, 1979; He et al., 2022; Ward et al., 2019). More carefully controlled water limitation can still induce stress responses without causing severe drought effects. The vegetative stage is particularly critical in the rice life cycle, typically requiring irrigated conditions to establish robust growth. However, there remains a significant gap in understanding how rice responds to moderate water limitation during this stage, especially regarding the integration of stomatal and non-stomatal adaptations and their combined effects on grain yield and quality.

The current study addresses this gap by examining the physiological and biochemical responses of Australian commercial elite rice lines to limited water stress during the seedling, vegetative and seed filling stages. By maintaining water at 60-65% field capacity rather than inducing severe drought, we sought to identify rice varieties that display effective water conservation strategies without compromising productivity. The specific objectives were to characterise stomatal and non-stomatal leaf traits associated with WUE in diverse Australian commercial rice varieties under limited water conditions; identify varieties with superior water conservation traits; and determine whether limited water stress during seedling through to grain filling stages affects grain quality parameters including milling efficiency and nutritional composition. This integrated approach provides valuable insights for pre-breeding efforts aimed at developing rice varieties that maintain productivity under water-limited conditions.

## Materials and methods

### Rice Germplasm

Twenty-one rice lines provided by the Department of Primary Industries, New South Wales, Australia, were evaluated. These included 18 Australian temperate *japonica* commercial rice lines, two *indica* rice varieties for comparison (Pokkali and Purple), and Moroberekan; a West African tropical *japonica* cultivar (Inukai et al., 1996) that exhibits drought- and rice blast-resistance (Grondin et al., 2018), as the positive control. The rice varieties represented different grain types including medium grain, long grain, short grain, fragrant, and arborio (Table S1).

### Soil field capacity (FC) determination

A soil dry-down curve was developed for the potting media (70% composted pine bark 0-5 mm, 30% coco peat, pH 6.35, EC = 650 ppm, with 3 g/L Osmocote Exact 3-4M [19-9-10 + 2MgO + TE, ICL Specialty Fertilizers], 2 g/L Osmocote Exact 5-6M [15-9-12 + 2MgO + TE]) (Vinarao et al., 2021) to determine the FC for the limited water experiment. Four soil pots (square-shaped, height 17 cm, width 7 cm) containing five rice plants each (10-week-old, different varieties) were saturated with water and left in the glasshouse (average temperature: 27°C, relative humidity: 50%) without adding water. The weights of the pots and soil moisture content (MC) were measured (Wireless soil moisture sensor, Ciderhouse Tech, AU) daily, and the plants’ drought response characteristics were observed throughout the experiment. Using early drought response such as slight leaf folding, drooping and leaf rolling, the wilting point was identified in the curve, and the midpoint from the beginning to the wilting point was selected as the FC to conduct the LW experiment.

### Glasshouse experiment

The experiment was conducted under two water treatments: ponded water (PW) and limited water (LW), in a glasshouse under controlled conditions at Swinburne University of Technology, Wantirna campus, Australia, during the summer of 2023. To mimic the rice-growing area in Australia, 16 hours of daylight at 25-30°C, 8 hours of night-time at 15-20°C, 50% average relative humidity, and light intensity of 30,000 - 35,000 lux were maintained in the glasshouse.

All seeds were primed at 40°C for four days and dehulled before planting. Pots with the same weight were prepared and placed in the water ponds in the glasshouse two days before seed planting to settle the potting mix. Four seeds were planted in each pot, and all other plants were removed at the three-leaf stage, leaving one plant per pot. Eight pots were prepared for each rice genotype, with four replicates conducted per variety under each water condition (PW: pots 1-4 and LW: pots 5-8). The experiment was conducted using a complete randomised block design.

When the plants in the LW plants pond reached the three- to five-leaf stage, water was removed from the LW plants. The MC determined through the dry-down curve (25-28%) was maintained throughout the entire vegetative growth stage (until panicle initiation). At this MC, the FC of the potting mix was 60-65% of the original FC. At panicle initiation, pots were re-irrigated until seed maturation.

### Data collection

#### Stomatal conductance (g_s_)

An SC-1 Leaf porometer equipped with desiccants in the sensor head (METER group, USA) was used to measure g_s_. The readings were generated on the lower surface of the rice leaves, with measurements taken from the middle, wider part of the fully expanded second mature leaves of individually grown plants.

#### Stomatal density

Imprints of the vegetative leaf epidermis were taken from the middle wider part of the fully expanded second mature leaves. A thin layer of fast-drying clear nail polish was applied and air-dried for about 10 minutes. Dry nail polish was peeled off using clear tape and placed on a microscope slide (Cowling et al., 2021; Pathoumthong et al., 2022; Wu & Zhao, 2017). Imprints of the abaxial (basal) and adaxial (upper) leaf surfaces were taken, and stomata were observed in a 600 μm × 450 μm region area through an EVOS 5000 Microscope (Thermo Fisher Scientific) at 200× magnification and converted into the number of stomata/mm^2^.

#### Cuticular and epicuticular wax (CEW) and flavonols

Attenuated Total Reflectance - Fourier Transform Infrared (ATR-FTIR) analysis (Nicolet iS5 Spectrometer, Thermo Fisher Scientific, Waltham, MA, USA) was used to analyse and semi-quantify the CEW and flavonols on fresh rice leaves (Sharma and Uttam, 2016; Willick et al., 2018; Yun et al., 2024). Spectra were measured in the 400 to 4000 cm^−1^ range using OMNIC software, with each spectrum representing the average of 32 scans at a resolution of 4 cm^−1^. Fresh leaves were dried in glassine bags for five days without applying pressure that could damage the epicuticle crystal structure. Dried samples were then sputter-coated with gold, and the structure was then observed using a scanning electron microscope (SEM) to compare changes between PW and LW leaves.

#### Photosynthetic parameters

The MultispeQ V 2.0 device (PHOTOSYNQ INC, USA) was used to measure the photosynthetic parameters of the leaves. Measurements included relative chlorophyll content, leaf temperature, chlorophyll fluorescence, quantum yield of photosystem II (ΦPSII), maximal quantum efficiency of PSII (Fv/Fm), total non-photochemical quenching (ΦNPQt), fraction of light dedicated to non-photochemical quenching (ΦNPQ), and fraction of light lost via non-regulated photosynthesis inhibitor processes (ΦNO).

#### Yield and milling quality of seed

Thousand grain weight (TGW) was determined by randomly selecting a representative sample of fully filled grains from each variety of PW and LW plants. The total mass of 1,000 grains was recorded. Paddy rice samples were dehulled using a laboratory-scale rice husker (TR-260 Automatic Rice Husker, Kett, Japan) to obtain brown rice, and brown rice yield (BRY) was calculated as the percentage of brown rice weight relative to the initial paddy weight. The brown rice was polished using a laboratory polisher (PEARLEST TP-3000 Grain Polisher, Kett, Japan) to obtain milled rice, and milling rice yield (MRY) was expressed as the percentage of milled rice weight relative to the initial paddy weight. Milling and polishing were performed for 60 seconds for 10 g of each sample. Head rice recovery (HRR) was determined by separating intact whole grains (≥75% of full grain length) from broken grains after milling and calculating the percentage of head rice weight relative to the initial paddy weight.

#### Nutritional quality of seeds

The same ATR-FTIR instrument used to detect CEW was employed for the surface analysis of the resulting seeds after dehulling, with each spectrum representing the average of 64 scans at a resolution of 4 cm^−1^.

#### Statistical analysis

Data visualisation was performed using GraphPad Prism version 10.0.1 and Python in Jupyter Notebook. Student’s t-test was conducted in GraphPad Prism to assess the significance of differences between the two water treatments, while a two-way ANOVA was used to analyse the effects of variety, water treatment, and their interaction. Statistical significance was set at p < 0.05. For each measurement, data represent the mean ± standard error of the mean, with n = 21 varieties, each with 4 biological replicates and appropriate technical replicates depending on the parameter being measured.

## Results

### Soil dry-down curve

Leaf rolling is a classic drought response in rice. Leaf rolling became evident in most plants on Day 7, as shown in Supplementary Figure S1. We thus used moisture content on Day 4 (25-28%) as a guide to maintaining limited water but not drought throughout the vegetative stage of the LW plants. At that moisture content, FC of the potting mix was ∼60-65% of the original FC.

### Stomatal Leaf Traits

#### Stomatal conductance (g_s_) and stomatal density

Most of the plants subjected to LW conditions exhibited reduced g_s_ than those under PW conditions during the vegetative stage at week 12, indicating a water-conserving response in the LW treated plants (Fig. 1A). The t-test revealed no significant difference in g_s_ between japonica and indica varieties under PW conditions (p = 0.1239), whereas a significant difference was observed under LW conditions (p < 0.0001). Generally, indica rice varieties tend to have higher stomatal density and g_s_ than japonica varieties (Shimoda & Maruyama, 2014; Zhang et al., 2022). Consistent with this, the indica varieties Pokkali and Purple exhibited higher g_s_ than most japonica varieties under ponded conditions. However, japonica varieties such as Amaroo, Bogan, Doongara, and Goolarah showed higher g_s_ than the indica varieties, with Bogan displaying the highest g_s_. Under limited water conditions, indica varieties demonstrated higher g_s_ than japonica, indicating that japonica varieties may have better water conservation abilities. Notably, Koshihikari, Reiziq, and Sherpa exhibited reduced sensitivity in gas exchange regulation under limited water stress. The adaxial axis reported reduced stomata than the abaxial axis for all varieties. Plant leaves under LW conditions exhibited reduced stomatal densities on the abaxial surface than those under PW conditions during the vegetative stage at week 12, suggesting a water-conserving adaptation (Fig. 1B). Doongara, Illabong, Lemont, Paragon, and Sherpa showed less change in stomatal density under water stress. Indica varieties had higher stomatal densities under both water treatments and experienced more pronounced reductions under LW conditions than most japonica varieties. Bogan displayed the greatest reduction, indicating a strong response to water stress. The t-test revealed a significant difference in stomatal density between japonica and indica varieties under both PW conditions (p < 0.0001) and LW conditions (p = 0.0003).

**Fig. 1.**
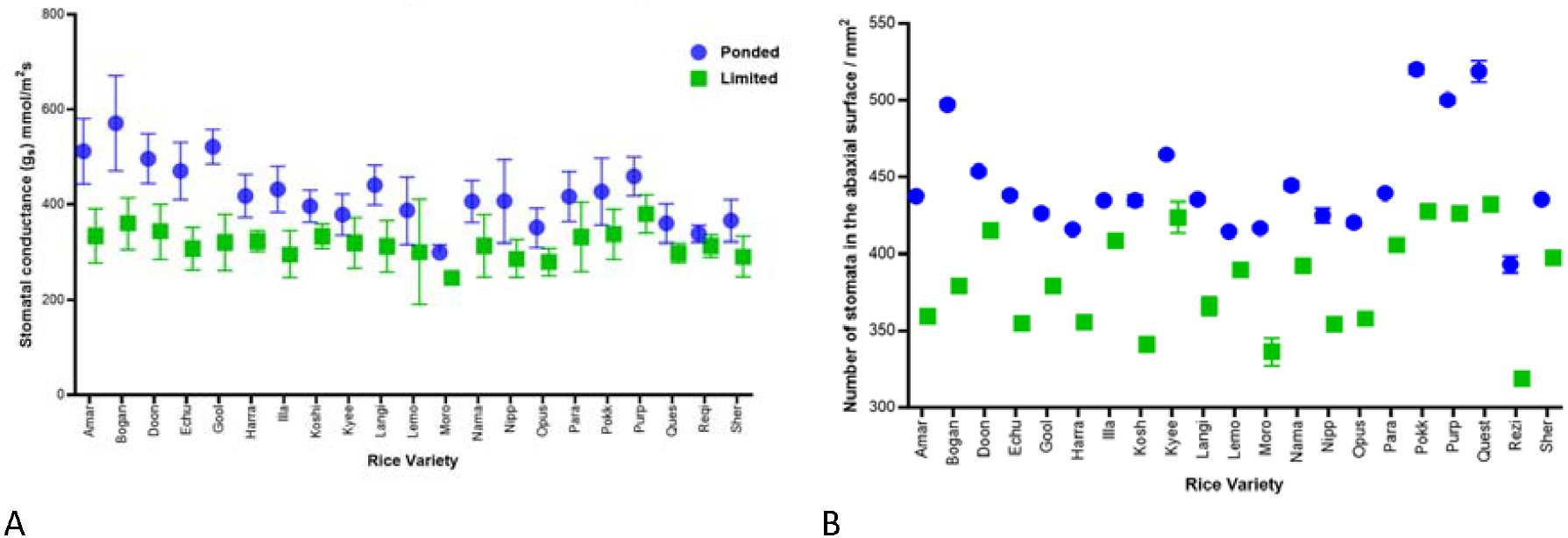
Stomatal traits of rice varieties under ponded water (PW) and limited water (LW) conditions at week 12. (A) Stomatal conductance (gs) showing reduced values under LW treatment (p < 0.0001, t-test; p < 0.001, two-way ANOVA). (B) Stomatal density on the abaxial leaf surface showing decreased values under LW conditions (p < 0.001, t-test and two-way ANOVA). Data represent mean ± SEM, n = 21 varieties with 4 biological replicates per variety per treatment and 12 technical replicates.

#### Non-stomatal leaf traits

Cuticular and epicuticular wax (CEW) **and Flavonols**. In the ATR-FTIR spectra (Supplementary Fig. S2) of fresh rice leaves, the peaks related to deposition of CEW were found in the spectral region 2800-3000 cm^−1^ due to symmetric and asymmetric stretching vibrations of the methyl and methylene groups, primarily caused by lipids (Phansak et al., 2021; Sharma & Uttam, 2016). Several peaks related to flavonols were identified, especially in the fingerprint area of the spectrum: 1125-1140 cm^−1^ (C-H bending ring vibration), 1205-1225 cm^−1^ (C=C ring stretch), 1270-1310 cm^−1^ (C=C stretching and OH bending vibrations), 1435-1475 cm^−1^ (ring stretch and OH bending vibrations), and 1605-1620 cm^−1^ (C=O and C2=C3 stretching vibration) (Herath et al., 2024; Krysa et al., 2022).

Figures 2 and 3 show a time series analysis of CEW deposition and flavonol formation on the leaves derived from the ATR-FTIR spectrum. At Week 3 (Wk3), all plants were under PW conditions just before starting the LW experiment, resulting in no significant difference between the PW and LW readings. During the water stress period, at weeks 10 (Wk10) and 12 (Wk12), the LW leaves had a higher amount of CEW and flavonols, and they achieved the optimum CEW deposition at an increasing rate. The salt tolerant Cv Pokkali (Tiwari et al., 2023) and stress resistant Cv Sherpa (Vinarao et al., 2023) produced a higher amount of CEW and flavonols during the water stress. After re-irrigating the plants, there was no significant difference in CEW between the readings, but the amount of flavonols showed a significant difference.

**Fig. 2.**
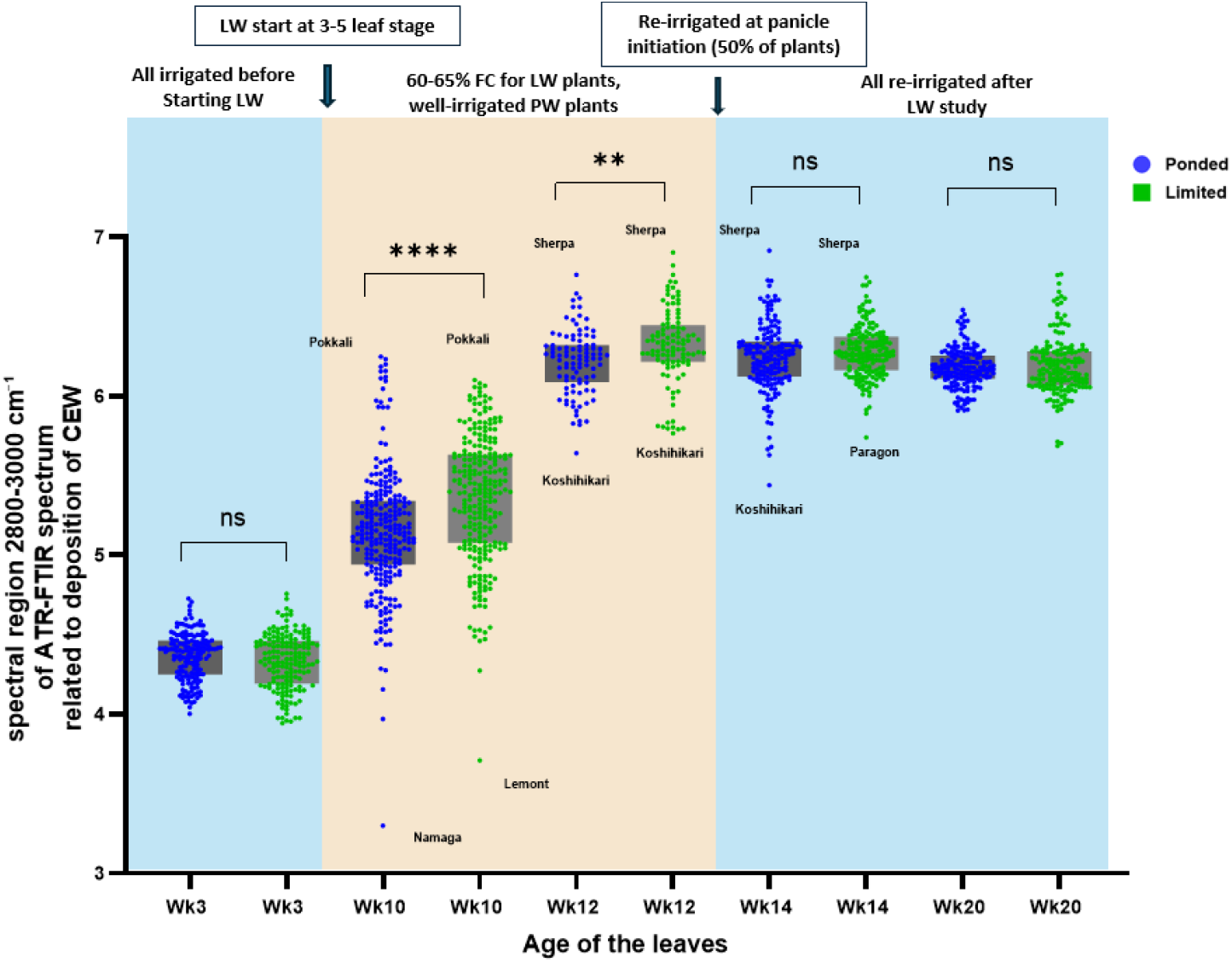
Time-course of cuticular and epicuticular wax (CEW) deposition on rice leaf surfaces under ponded water (PW) and limited water (LW) conditions. The brown shaded area indicates the experimental period when LW treatment was applied. LW leaves showed significantly higher CEW deposition during water stress at weeks 10 and 12, with differences diminishing after re-irrigation. Data represent mean ± SEM, n = 21 varieties with 4 biological replicates per variety per treatment and 12 technical replicates.

**Fig. 3.**
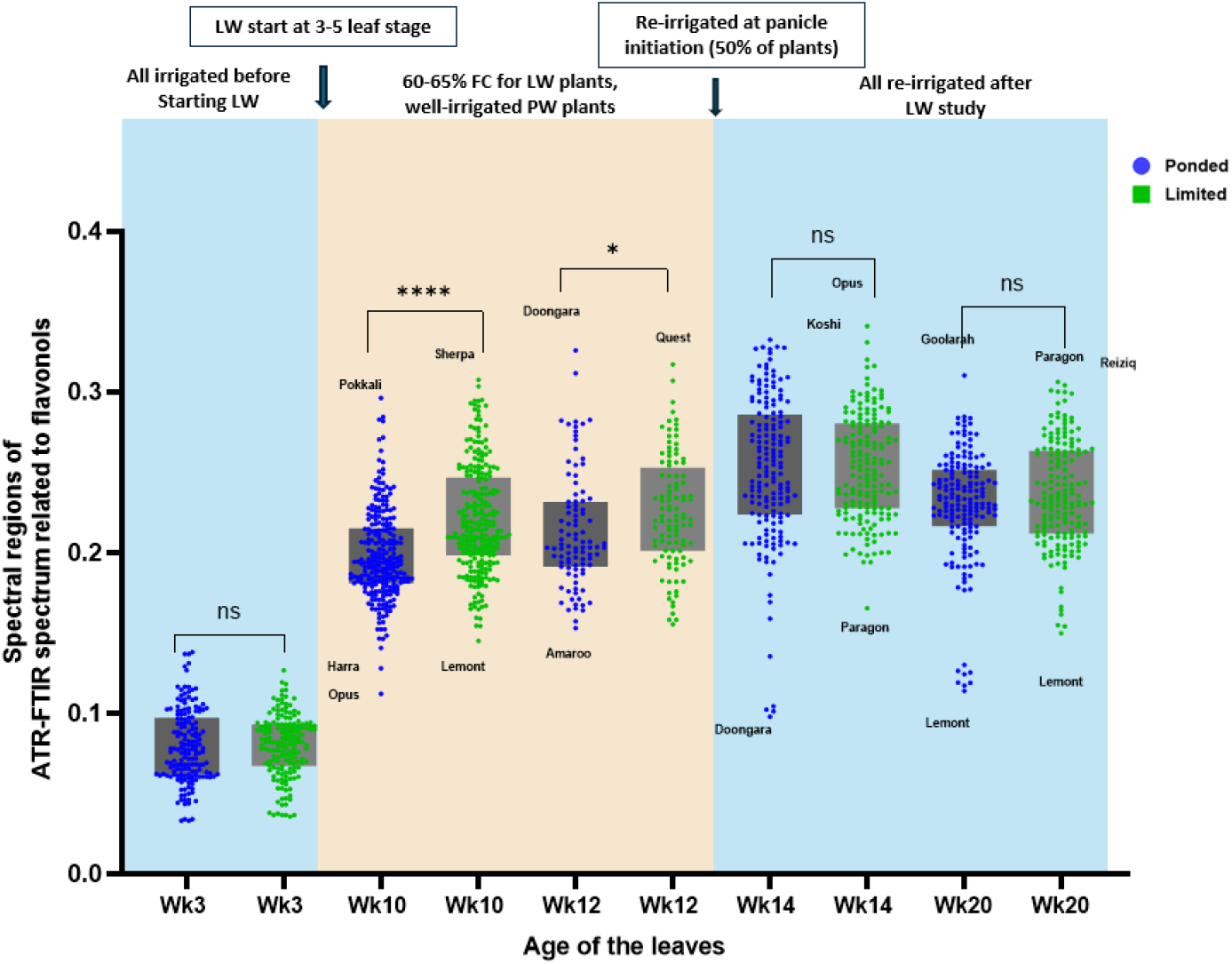
Time-course of flavonol content on rice leaf surfaces under ponded water (PW) and limited water (LW) conditions. The brown shaded area indicates the experimental period when LW treatment was applied. LW leaves exhibited significantly higher flavonol content during water stress, with differences persisting after re-irrigation. Data represent mean ± SEM, n = 21 varieties with 4 biological replicates per variety per treatment and 12 technical replicates.

#### SEM observation

Both the LW and PW leaves had the same crystal-like epicuticular wax structures, but the LW leaves exhibited larger and more elevated papillae on their surface. It was observed that unlike the papillae on the PW leaves, the larger papillae on the LW leaves cover the stomata openings well (Fig. 4).

**Fig. 4.**
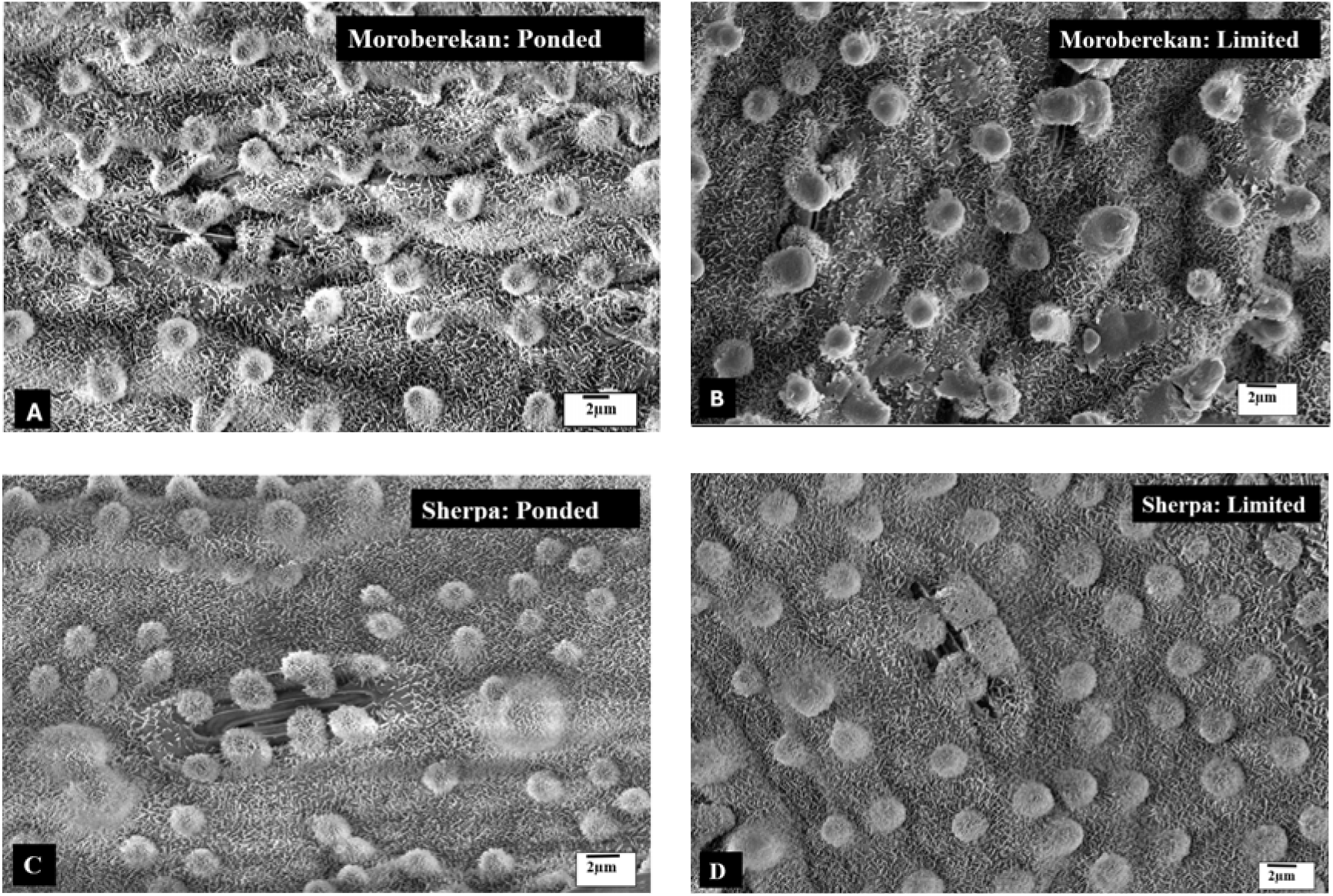
Scanning electron microscopy (SEM) images showing epicuticular wax structures on leaf surfaces of Moroberekan and Sherpa rice varieties. (A, C) PW leaf surface showing normal epicuticular wax crystals and papillae. (B, D) LW leaf surface displaying larger and more elevated papillae that cover stomatal openings. Scale bars represent 2 µm. Images shown are representative of 4 biological replicates per treatment with 3 technical replicates each.

#### Photosynthetic parameters

LW plants showed reduced relative chlorophyll content, ΦPSII, and Fv/Fm than PW plants, indicating reduced photosynthetic efficiency as they had limitations in taking in CO_2_ through stomata for photosynthesis (Fig. 5A, C, D). Higher leaf temperatures in LW conditions were observed because of reduced transpiration (Fig. 5B). Higher ΦNPQ values were observed in LW plants (Fig. 5E). ΦNPQ measures the proportion of absorbed light energy that a plant dissipates as heat, acting as a protective mechanism to prevent damage from excess light. The increased ΦNPQ in LW plants reflects their higher need to dissipate excess light energy due to reduced photosynthetic capacity under water stress. Elevated ΦNO values generally indicate that a plant is experiencing significant stress (Fig. 5G). These higher values are often associated with increased non-photochemical energy dissipation as a response to stress conditions.

**Fig. 5.**
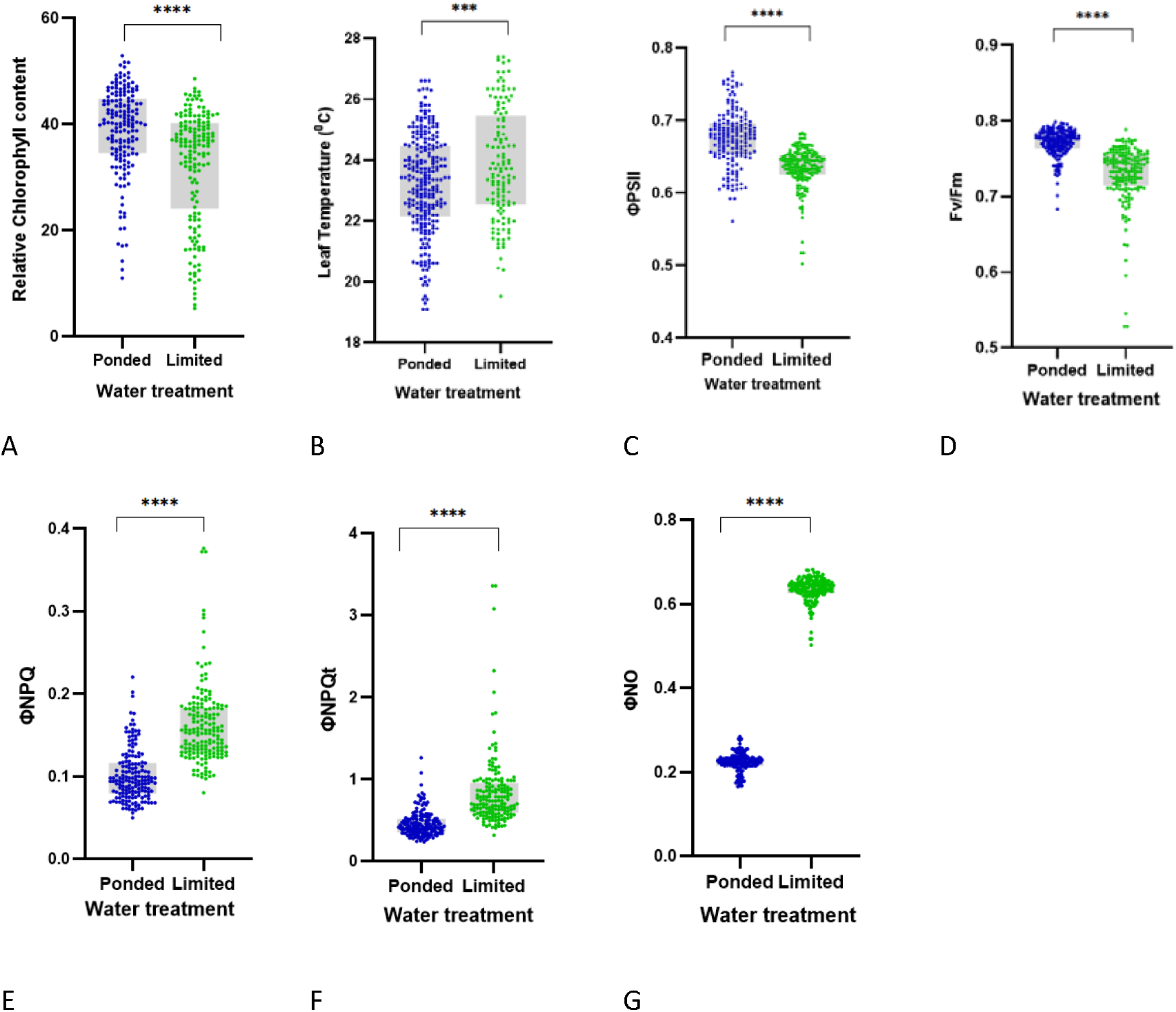
Photosynthetic parameters of rice varieties under ponded water (PW) and limited water (LW) conditions at week 12. (A) Relative chlorophyll content, (B) leaf temperature, (C) quantum yield of photosystem II (ΦPSII), (D) maximum quantum efficiency of PSII (Fv/Fm), (E) fraction of light dedicated to non-photochemical quenching (ΦNPQ), (F) total non-photochemical quenching (ΦNPQt), and (G) fraction of light lost via non-regulated photosynthesis inhibitor processes (ΦNO). LW plants showed significant decreases in photosynthetic efficiency parameters and increases in photoprotective mechanisms. Data represent mean ± SEM, n = 21 varieties with 4 biological replicates per variety per treatment and 12 technical replicates.

**Fig. 6.**
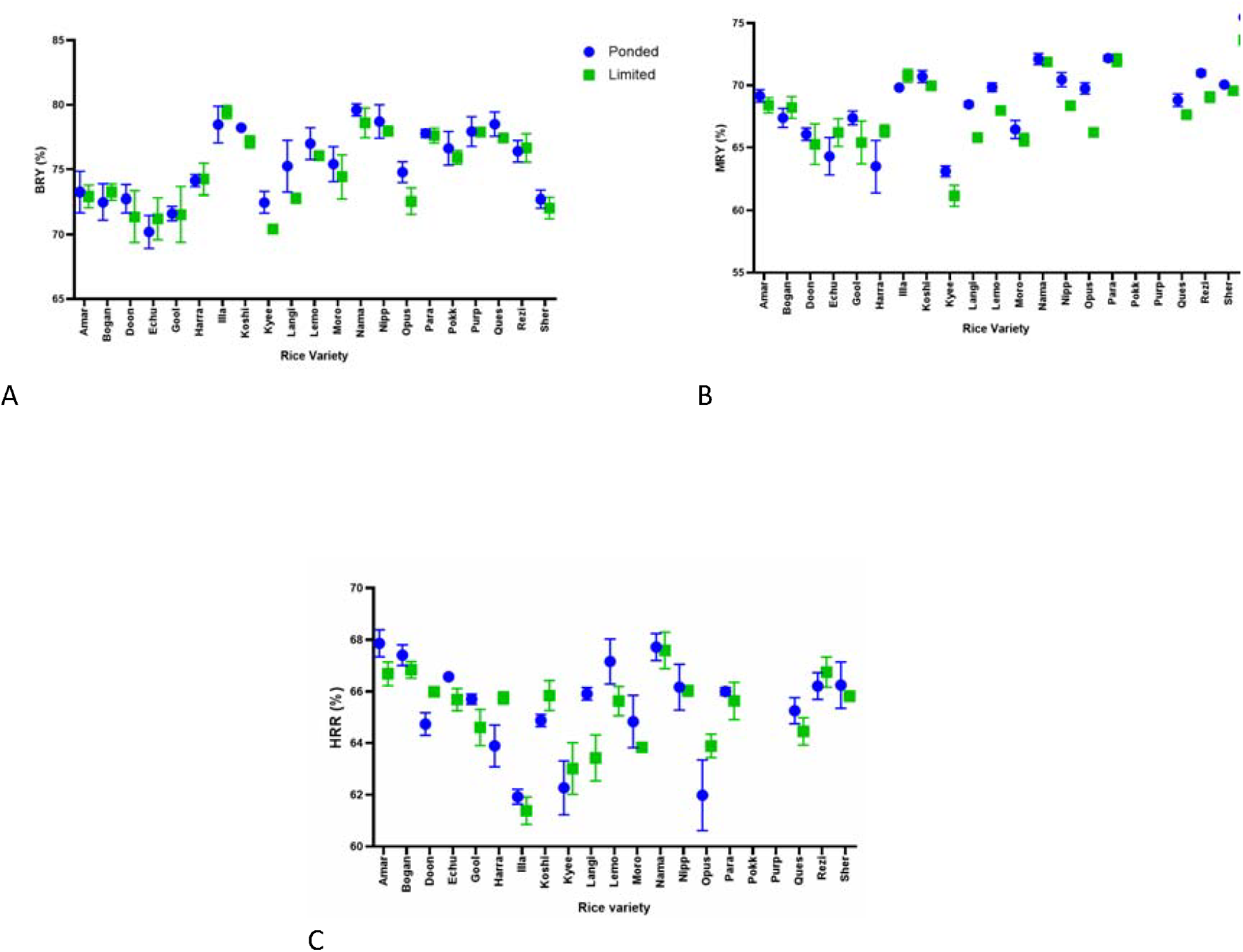
Milling recovery traits of rice varieties grown under PW and LW conditions. (A) BRY was significantly influenced by rice variety (p < 0.0001) and water treatment (p = 0.0019), but showed no significant interaction effect (p = 0.2481). (B) MRY was significantly influenced by rice variety (p < 0.0001) and water treatment (p = 0.0019), with a significant interaction effect (p = 0.0062). (C) HRR was significantly influenced by rice variety (p < 0.0001) and interaction effect (p = 0.0062), but water treatment (p = 0.3119) with no significant effect. Unpaired t-tests showed no significant overall differences between treatments for BRY (p = 0.4828), MRY (p = 0.3834) or HRR (p = 0.7124). Data represent mean ± SEM, n = 21 varieties with 4 biological replicates per variety per treatment and 4 technical replicates.

A scatter plot with a regression line was used to visualise the relationship between ΦPSII, ΦNPQ, and Fv/Fm under PW and LW conditions, providing insights into the degree of correlation and variability among rice varieties in response to water stress (Supplementary Fig. S3). Opus and Paragon have relatively low ΦPSII in both conditions. Langi, Kyeema, and Moroberekan show higher ΦPSII under both conditions, suggesting better adaptability. The red shaded region represents the 95% confidence interval of the regression line. Varieties above the red line perform better under LW, and varieties below the line may be more sensitive to water limitations. Higher ΦNPQ values in LW leaves indicate that plants are under greater stress, reflecting the plant’s need to dissipate more excess light energy to protect itself from damage. Paragon represents a particularly sensitive response to water stress, showing a much higher ΦNPQ under LW conditions and significant damage to its photosynthetic machinery, leading to a low Fv/Fm. Moroberekan is a variety reported to be more tolerant to drought stress, maintaining its photosynthetic efficiency relatively well under stress. Sherpa appears to have a moderate response to water stress in terms of ΦNPQ, while Purple might have a slightly enhanced ability to activate non-photochemical quenching under stress. According to the plots, Opus might also be more sensitive to water stress.

#### Yield and milling quality of seeds

The important yield trait TGW (Supplementary Fig. S4) varied significantly among rice varieties and between water treatments, confirming the influence of both genetic and environmental factors. However, the difference in TGW between PW and LW conditions was not significant, suggesting that LW during the vegetative stage did not negatively impact grain weight. Although some long-grain varieties, such as Doongara, Kyeema, Langi, and Lemont, showed reduced TGW under limited water, other varieties maintained stable TGW across treatments. The non-significant interaction (p = 0.0636) indicates that the response of varieties to water scarcity was consistent. These results suggest that limited water stress during the vegetative stage may allow for water savings without compromising grain weight, which could help maintain yield.

Milling recovery traits, including brown rice yield (BRY) and milling rice yield (MRY), were significantly influenced by both rice variety and water treatment, but the interaction effects differed between the two traits. BRY did not show a significant interaction effect, suggesting a consistent response to water availability across varieties. In contrast, MRY exhibited a significant interaction, indicating that the impact of water treatment on milling recovery varied by variety. The effects of water treatments on the outcome of the experiment are influenced by the rice variety used, highlighting a significant interaction between these two factors. However, rice variety itself plays a more dominant role in determining the overall outcome, with its contribution to the variation being considerably higher than that of water treatments. Despite these differences, the unpaired t-tests for BRY and MRY showed no significant overall difference between PW and LW treatments, suggesting that LW during the vegetative stage did not negatively affect milling recovery. Head rice recovery (HRR) was significantly influenced by rice variety and showed a significant interaction effect between variety and water treatment, indicating variety-specific responses. However, the absence of a significant overall effect of water treatment suggests that moderate drought during the vegetative stage did not adversely affect grain processing quality, as measured by HRR. Since HRR is closely associated with market value and consumer quality perception, these findings highlight the potential for using LW during the vegetative stage without compromising commercially important quality traits. Only BRY was determined for two indica varieties, as they are commonly marketed as brown or minimally milled rice due to their colour. Pokkali is light brown and purple is a pigmented rice with high anthocyanin.

#### Nutritional Quality of Seeds

The peak area that was used for definition of protein content were are found in the spectral region 1500-1570 cm^−1^ associated with N-H bending and C-N stretching vibrations and 1620-1640 cm^−1^ primarily due to C=O stretching vibrations. Peaks related to lipids are found at 1740-1730 cm^−1^ associated with ester carbonyl groups in triglycerides and phospholipids, and C-H stretching around 2850-2950 cm^−1^, indicative of aliphatic chains in lipids. A peak at 1031 cm^−1^ was described to be associated with the C-O stretching vibrations of polysaccharides, including starch and other carbohydrates (Ishikawa et al., 2021; Siregar et al., 2024) (Supplementary Fig. S5).

As shown in Fig. 7, the protein content derived from the spectral reading suggest it was significantly greater in plants subjected to LW treatments, with both water treatment and variety having substantial effects. Lipid content did not differ significantly between treatments, although a variety-specific interaction was observed. Starch content did not change between water treatments, or with variety. Results suggest that LW stress can affect protein content but has a minimal impact on lipids and starch in rice seeds (Supplementary Fig. S6).

**Fig. 7.**
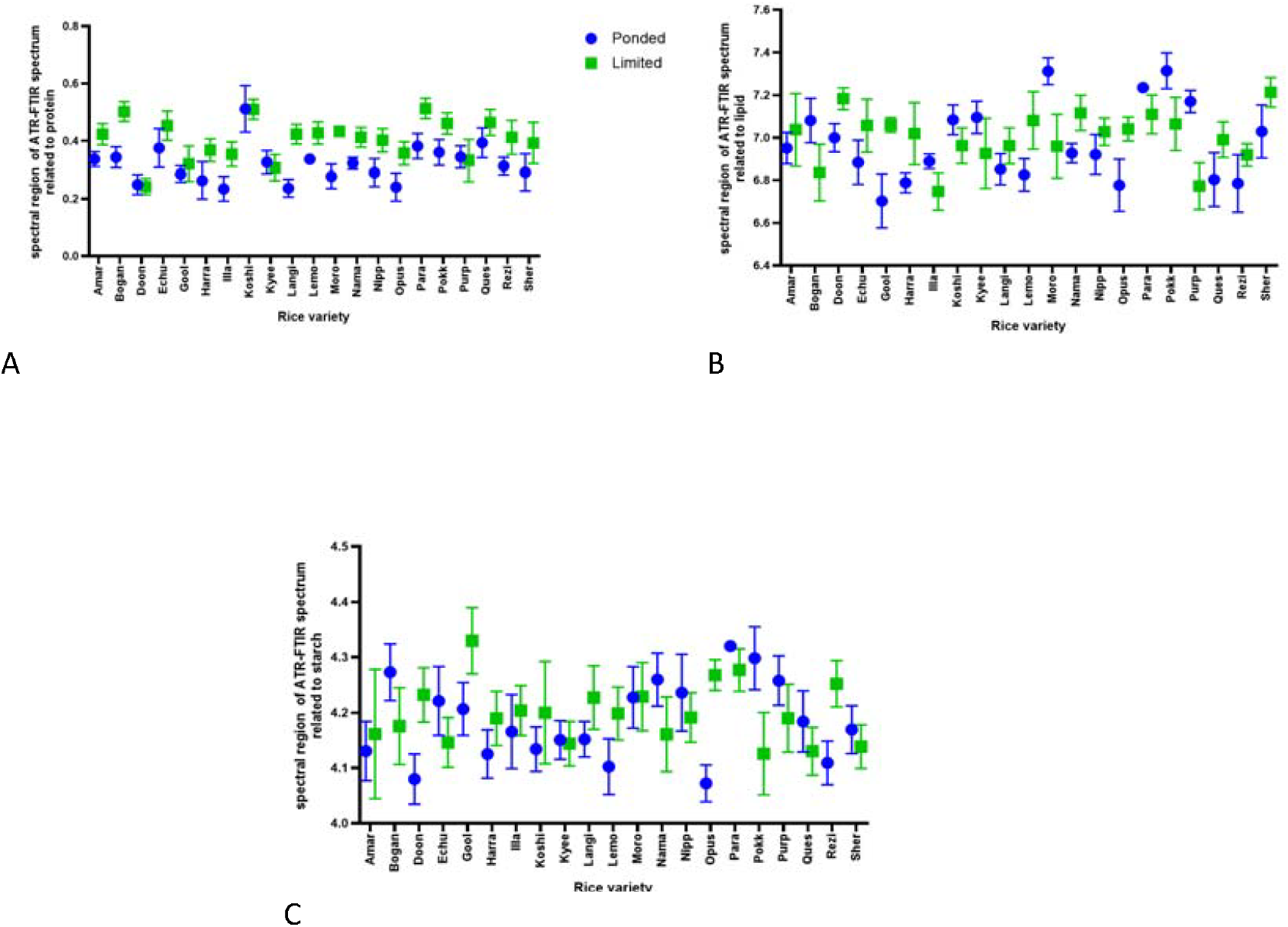
Nutritional composition of rice seeds from plants grown under ponded water (PW) and limited water (LW) conditions. (A) Protein content was significantly higher in LW plants (p = 0.0446, t-test; effects of water treatment p < 0.0001 and variety p < 0.0001, two-way ANOVA). (B) Lipid content showed no significant difference between treatments (p = 0.8983, t-test), but a significant interaction between water treatment and variety (p = 0.0024). (C) Starch content showed no significant differences between treatments (p = 0.2022, t-test) or effects of water treatment or variety (p > 0.05 for both, two-way ANOVA). Data represent mean ± SEM, n = 21 varieties with 4 biological replicates per variety per treatment and 12 technical replicates.

## Discussion

Rice can cope with water scarcity at different levels. Stress resistant plants can evade the stress condition by changing plant architectural traits or on the physiological and molecular level. Hereby also the stage of development as well as the severity and duration of applied stress is of importance. Inducing drought at different stages of the rice life cycle for shorter or longer periods (Baisakh et al., 2020; Kamoshita et al., 2015; Lafitte et al., 2007) can negatively impact both yield and quality of rice (Haque et al., 2016; Khan et al., 2021; Pitaloka et al., 2021; Yang et al., 2019). Breeding to improve the WUE of rice under LW conditions is advisable to help maintain productivity and grain quality. The FC used for this study (60-65% of the original FC of the potting mix) is almost identical to that used by (Reddy et al., 2020) to create LW stress (albeit only from 45 to 80 days after sowing, not throughout the vegetative stage). Most previous drought studies have been conducted at < 50% FC from the original FC of the soil (Lafitte et al., 2007; Pitaloka et al., 2021; Praba et al., 2009; Siopongco et al., 2006).

Previous drought studies have mostly reported significantly reduced g_s_ in water-stressed plants. In the present study, however, LW-stressed plants maintained greater g_s_ and a more efficient gas exchange strategy allowing sufficient carbon assimilation for photosynthesis while minimizing excessive water loss which may contribute to improved yield. Farooq et al. (2010) mentioned that drought-stressed plants show gs of 30-65 mmol/m^2^s and well-watered plants show 287-395 mmol/m^2^s, while Ouyang et al. (2017) showed that drought-stressed rice plants show much reduced g_s_ (less than 150 mmol/m^2^s) than mild drought and well-watered plants. Dien et al. (2017) reported a significant decrease in g_s_ to below 100 mmol/m^2^/s after 10 days of drought, while well-watered plants maintained gs levels between 400 and 500 mmol/m^2^/s in both indica and japonica rice varieties. This included the variety “Moroberekan”, which was reported as drought stress resistant (Grondin et al., 2018) and therefore included as reference in this work.

Reduced number of stomata conserve water but also limit carbon assimilation, requiring careful management (Caine et al., 2019). Moderate drought conditions contribute to a gradual decrease in stomata numbers, while severe drought leads to a more significant reduction (Ilyas et al., 2021). In the current study, LW plants had stomatal densities of more than 300/mm^2^, higher than those typically seen in drought conditions, which is around 250/mm^2^ (Freeg et al., 2022), suggesting improved yields. Caine et al. (2019) found that increased CO_2_ and higher temperatures could enable lower stomatal density varieties to maintain sufficient gas exchange and achieve equivalent or improved yields, even with reduced photosynthesis. Thus, slightly reducing stomatal density may not negatively impact rice yields in the future with rising CO_2_ and temperatures. In addition to density, stomata in LW plant leaves were also found to be covered with larger and more elevated papillae, a structural adaptation that may further enhance water conservation by reducing transpiration directly at the stomatal pore.

Previous studies have shown that rice leaves produce more CEW under water stress (Haque et al., 1992; Srinivasan et al., 2008), but little is known about the role of wax papillae in managing stresses (Pitaloka et al., 2021). The larger papillae that cover the stomata, observed on the LW plant leaves in this study, might be a key adaptation that helps rice leaves conserve water and protect against various abiotic and maybe biotic stresses. On the other hand, flavonoids help plants cope with water stress by acting as antioxidants, scavenging reactive oxygen species (ROS), and protecting against UV damage. They also regulate stress-related hormones, maintain osmotic balance, and enhance root development, improving the overall drought tolerance of plant (Laoué et al., 2022; Shomali et al., 2022). The results of the current study show that flavonols play a promising role in stress tolerance in rice leaves. Moreover, the results of this study indicate that when LW leaves reach the optimal wax level at an increasing rate, they produce more flavonols than PW leaves.

The reduction in ΦPSII and the increase in NPQ observed in water-stressed plants in the current study are less pronounced than in previous drought experiments (Caine et al., 2019; Pitaloka et al., 2022), suggesting a potential for better yield under LW. These findings are consistent with those of Vijayaraghavareddy et al. (2022), who reported similar trends in ΦPSII and NPQ in a LW experiment at 60% FC.

Grain physical and milling quality traits are essential for assessing rice performance under different water conditions. TGW, along with milling quality parameters such as BRY, MRY, and HRR, determines grain development, processing efficiency, and market value. TGW reflects grain size, density, and the grain-filling process, influenced by genetic and environmental factors, particularly water availability. While primarily a yield-related trait, TGW also impacts milling efficiency by affecting whole grain recovery (Bao, 2019; Butardo & Sreenivasulu, 2019; Zuo et al., 2022).

Milling quality, a key factor in consumer preference and marketability, is evaluated through BRY, MRY, and HRR. BRY measures whole grain retention after dehulling, MRY represents total milled rice recovery, and HRR is critical for market value as it indicates the proportion of intact head rice (Bao, 2019; Juliano, 1993). The statistically non-significant difference in TGW between PW and LW conditions is a significant finding, underscoring the potential of limited water irrigation strategies to maintain TGW, especially in short and medium grain varieties. This result offers promising prospects for sustainable rice cultivation, demonstrating that water conservation through using limited water during the vegetative stage can be achieved without compromising yield quality. According to the results of this study, which shows no significant overall difference between PW and LW treatments in BRY and MRY, rice can maintain stable milling performance under limited water conditions during plant growth, though variety-specific responses in MRY emphasise the importance of selecting resilient varieties for water-saving cultivation.

Although FTIR-ATR has been used to identify spectral regions associated with protein, lipid, and starch in rice (Herath et al., 2024), it is important to note that the technique provides semiquantitative analysis rather than absolute quantification. Nevertheless, the consistent spectral differences observed between PW and LW treatments suggest genuine compositional changes in response to water limitation. The increased protein content observed in rice grains under LW stress aligns with previous studies showing that moderate water stress can sometimes enhance grain protein concentration (He et al., 2022), possibly because of altered nitrogen metabolism and translocation under water-limited conditions.

The stability of lipid and starch content between water treatments is a positive finding for grain quality preservation, as these components significantly influence cooking and eating qualities of rice. The variety-specific interaction observed for lipid content suggests that genetic factors modulate the response of lipid metabolism to water limitation, which could be relevant for breeding programs aiming to develop varieties with stable grain quality across variable water conditions.

Among the tested varieties, Sherpa showed notable resilience to limited water stress, maintaining relatively stable physiological parameters while showing adaptive responses through increased CEW deposition. These characteristics, combined with stable grain yield and quality metrics, position Sherpa as a promising candidate for water-efficient rice production systems. The differences observed between japonica and indica varieties in their responses to limited water stress highlight the importance of subspecies-specific breeding approaches for improving WUE. To our knowledge, this is one of the few studies to comprehensively track the effect of leaf traits on the physiological response of rice throughout its entire life cycle, from seedling to grain production stage. Moreover, it uniquely establishes the connections between early vegetative stage physiological adaptations to water limitation and subsequent milling and grain quality traits, providing valuable insights for developing water-efficient rice varieties that maintain end-use quality.

## Conclusion

Maintaining limited water conditions throughout the vegetative stage resulted in less reduction in gas exchange, number of stomata, and photosynthetic efficiency, along with reduced overall plant stress compared to previous drought studies. This approach may contribute to improved yield, which cannot be expected under severe drought conditions. Among tested varieties in the current study, Sherpa is a strong candidate for breeding rice varieties with enhanced WUE due to its reduced sensitivity in gas exchange regulation, less sensitivity in changes to stomatal density and photosynthetic efficiency, and the highest cuticular and epicuticular wax deposition among the japonica varieties. The findings of this study demonstrate that limiting water during the vegetative stage can induce water-conserving traits without compromising yield or grain quality, providing valuable insights for developing water-efficient rice varieties for sustainable production in water-limited environments.

## Abbreviations

ATR-FTIR: attenuated total reflectance Fourier transform infrared spectroscopy
BRY: brown rice yield
Cv: cultivar
CEW: cuticular and epicuticular wax
FC: field capacity
Fv/Fm: maximal quantum efficiency of PSII Gs stomatal conductance
HRR: head rice recovery
LW: limited water
MC: moisture content
MRY: milling rice yield
PW: ponded water
SEM: scanning electron microscope
TGW: thousand grain weight
ΦNO: fraction of light lost via non-regulated photosynthesis inhibitor processes
ΦNPQ: fraction of light dedicated to non-photochemical quenching
ΦNPQt: total non-photochemical quenching
ΦPSII: quantum yield of photosystem II

## SUPPLEMENTARY FIGURE CAPTIONS

**Fig. S1.** Soil dry-down curve for the potting mix used in the experiment. Plants showed drought resistance response (leaf rolling) on day 7, indicating the wilting point. The moisture content at day 4 (25-28%) was selected for maintaining limited water conditions, corresponding to 60-65% of field capacity.

**Table S1**. List of rice varieties used in the experiment and their classification. The table includes 18 Australian temperate japonica commercial rice lines, two indica varieties (Pokkali and Purple), and Moroberekan (positive control), along with their subspecies classification and grain type.

## Acknowledgements

We would like to express our gratitude to the Department of Primary Industries, New South Wales, Australia, for providing the rice germplasm used in this study. We also thank AgriFutures Australia for providing the funding for this research project through Project No. PRP-013062 and PRO-016914.

## Author contributions

YF: conceptualisation, methodology, investigation, data curation, formal analysis, visualisation, writing - original draft; MA: conceptualisation, supervision, writing - review & editing; MK: supervision, writing - review & editing; VB: conceptualisation, resources, supervision, funding acquisition, project administration, writing - review & editing.

## Data availability

The data underlying this article will be shared on reasonable request to the corresponding author.

---

**Fig. S2.**
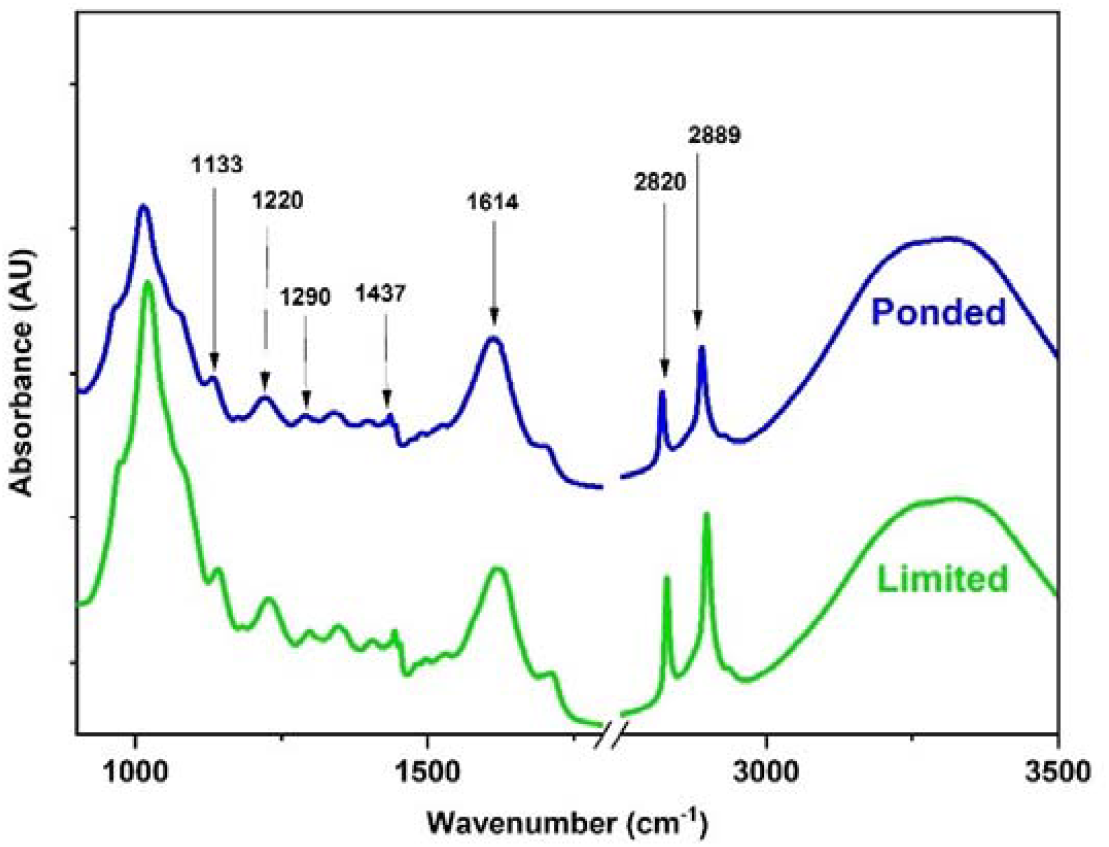
ATR-FTIR spectra of fresh rice leaves from ponded water (PW) and limited water (LW) plants. Key peaks identified include those for cuticular and epicuticular waxes (2800-3000 cm^−1^) and flavonols (1125-1140 cm^−1^, 1205-1225 cm^−1^, 1270-1310 cm^−1^, 1435-1475 cm^−1^, and 1605-1620 cm^−1^). Representative spectra shown are from Sherpa variety.

**Fig. S3.**
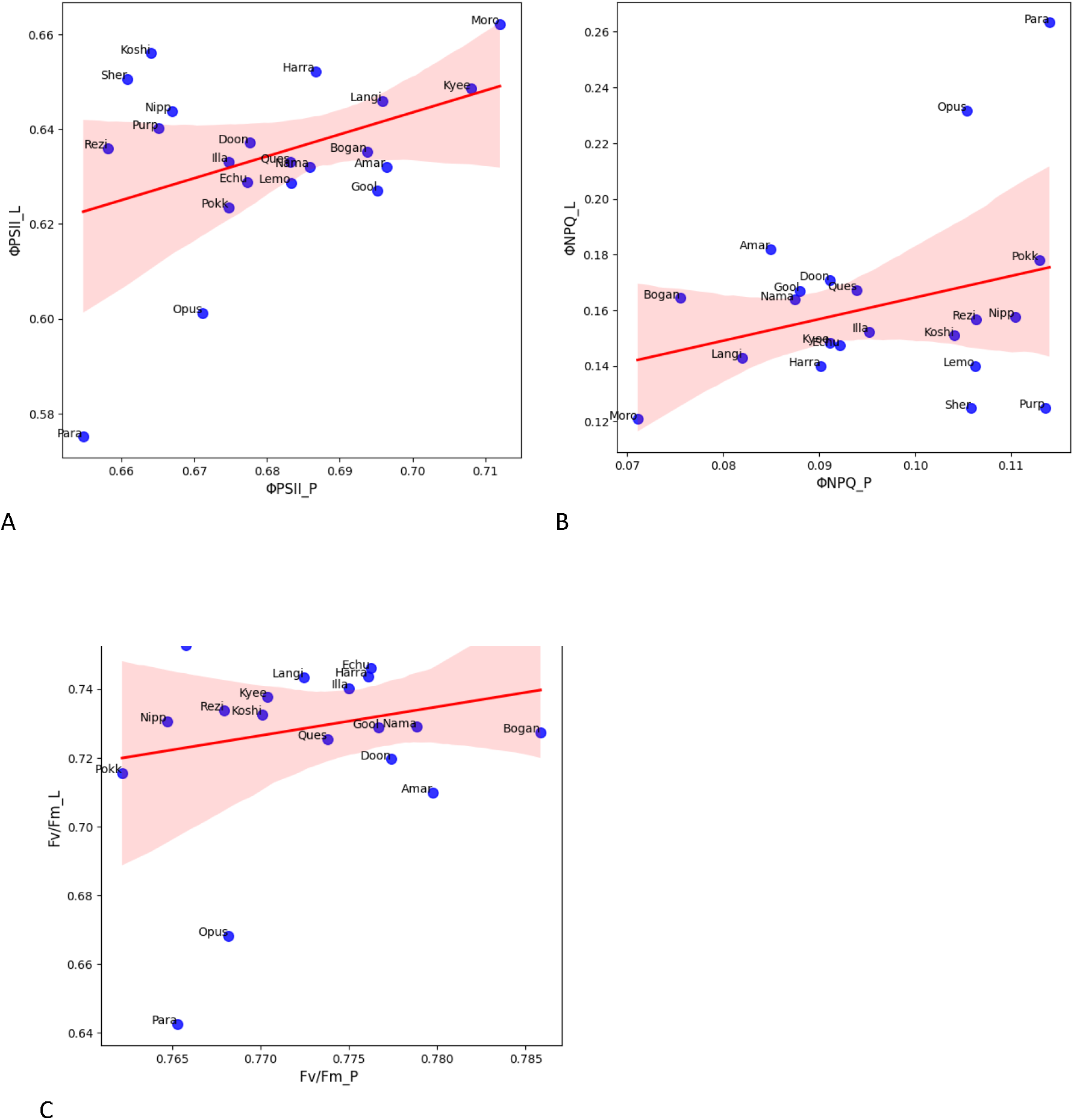
Relationships between photosynthetic parameters under ponded water (PW) and limited water (LW) conditions. (A) Quantum yield of photosystem II (ΦPSII), (B) fraction of light dedicated to non-photochemical quenching (ΦNPQ), and (C) maximum quantum efficiency of PSII (Fv/Fm). Each point represents a rice variety, with red regression lines and 95% confidence intervals (shaded areas). Varieties above the regression line show better performance under LW conditions. Data represent the average of 4 biological replicates per variety per treatment with 12 technical replicates.

**Fig. S4.**
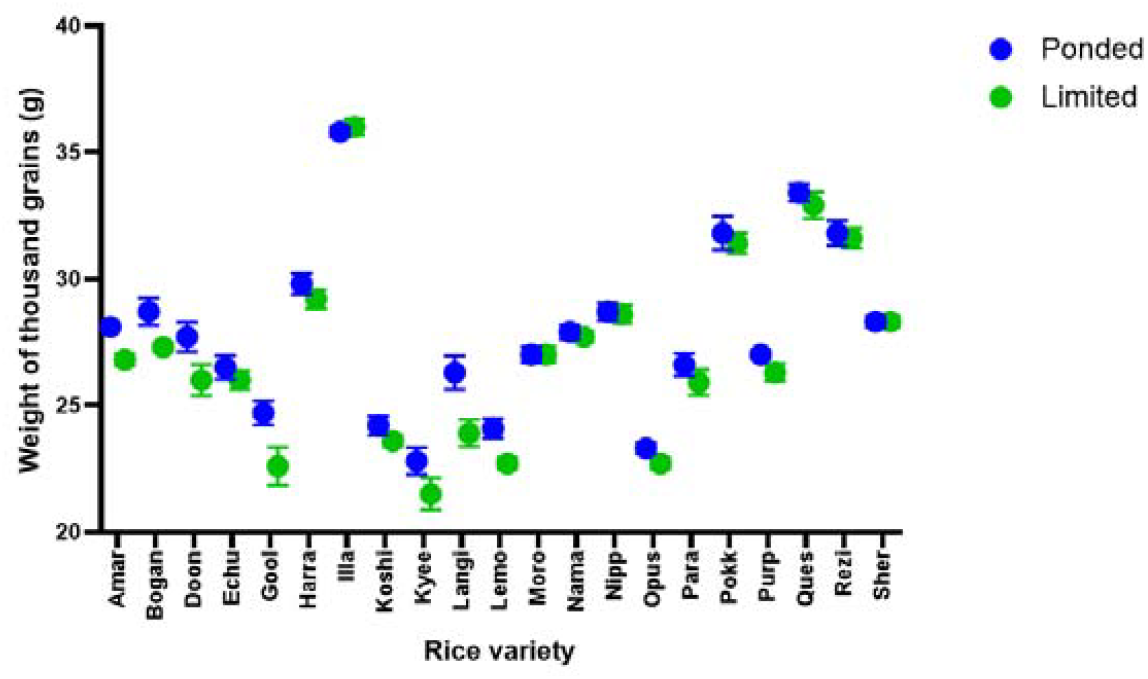
Thousand grain weight (TGW) of rice varieties grown under ponded water (PW) and limited water (LW) conditions. Significant effects of rice variety (p < 0.0001) and water treatment (p < 0.0001) were observed, but no significant overall difference between water treatments (p = 0.4752, t-test). Data represent mean ± SEM, n = 21 varieties with 4 biological replicates per variety per treatment and 4 technical replicates.

**Fig. S5.**
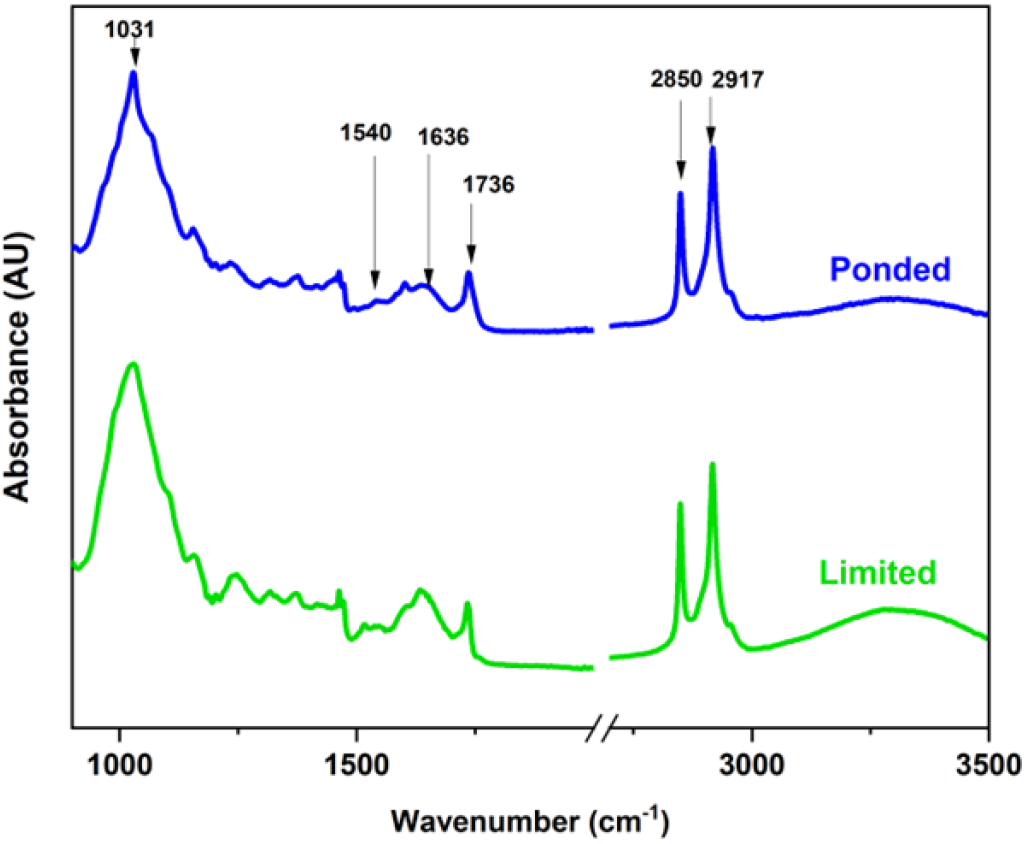
ATR-FTIR spectra of dehulled rice seeds from PW and LW plants. Key peaks identified include those for protein (1500-1570 cm^−1^, 1620-1640 cm^−1^), lipids (1730-1740 cm^−1^, 2850-2950 cm^−1^), and starch (1031 cm^−1^). Representative spectra shown are from Sherpa variety.

**Fig. S6.**
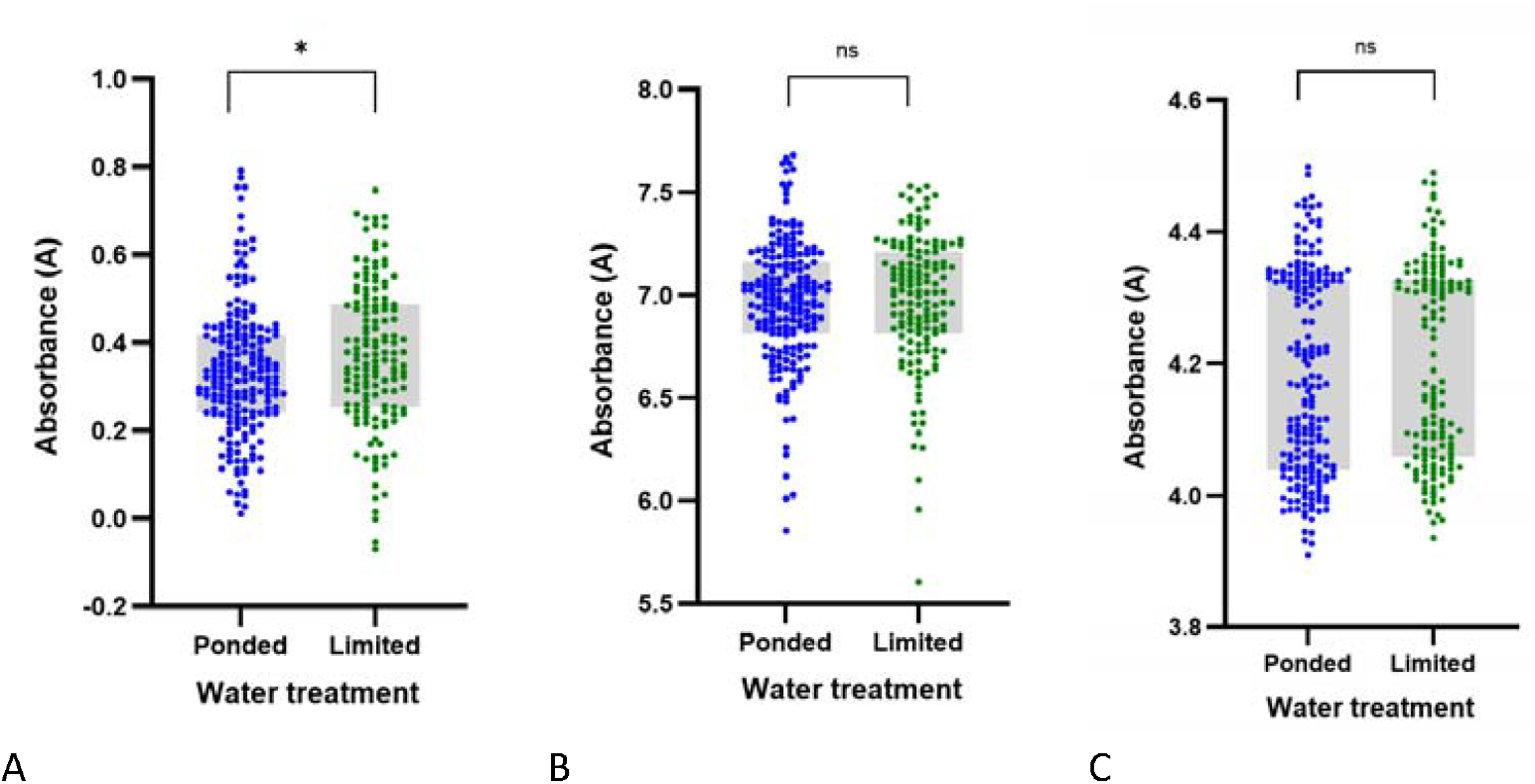
Comparison of nutritional components in rice seeds from ponded water (PW) and limited water (LW) plants across all varieties, showing the distribution of (A) protein, (B) lipid, and (C) starch content. Protein content showed significant increases under LW conditions, while lipid and starch content remained relatively stable.

